# Sphingolipid-driven interleaflet coupling orchestrates Rho-GTPase recruitment to nanodomains for signal activation in plants

**DOI:** 10.1101/2025.11.06.686946

**Authors:** Matheus Montrazi, Arthur Poitout, Camille Depenveiller, Vincent Bayle, Minoru Nagano, Adiilah Mamode Cassim, Marie-Dominique Jolivet, Jean-Bernard Fiche, Catherine Sarazin, Laetitia Fouillen, Françoise Simon-Plas, Jean-Marc Crowet, Yvon Jaillais, Sébastien Mongrand, Alexandre Martinière, Yohann Boutté

## Abstract

Biological membranes are both laterally heterogeneous and asymmetrical across leaflets, yet how this asymmetry contributes to signal transduction remains unclear. Here we show that sphingolipid-driven interleaflet coupling coordinates nanodomain organization and Rho-GTPase activation in plants. Using molecular dynamics simulations, super-resolution and single-molecule imaging, quantitative genetics, and biochemistry, we find that very long acyl chain (VLCFA)–containing sphingolipids in the outer leaflet interdigitate with phosphatidylserine (PS) in the inner leaflet, forming a vertical molecular bridge that organizes PS into nanodomains. This coupling promotes recruitment and activation of the Rho-GTPase ROP6 in response to auxin, whereas disruption of VLCFA synthesis or sphingolipid composition disperses PS and ROP6 nanodomains, impairing cytoskeletal reorganization and directional growth. Our findings reveal interleaflet coupling as a fundamental organizing principle linking membrane asymmetry to signaling, providing a conceptual framework for spatial and temporal control of signal transduction across eukaryotic membranes.

---

Cell signaling is a pivotal process that enables organisms to respond with precision to environmental and developmental *stimuli*. At the molecular level, biochemical information is transmitted through tightly regulated reactions in time and space. This is exemplified by the lateral organization of the receptors, co-receptors and their downstream effectors within nanoscale signaling platforms, also designated as nanodomains^1,2^ at the plasma membrane (PM). This spatial organization within the plane of the membrane is crucial for initiation of cell signaling but also to generate signal specificity^3,4^. While lipid nanodomains can be studied using *in vitro* systems, experimental evidence on how these nanodomains are formed and maintained *in vivo* remains scarce^5–9^. It has recently been demonstrated by means of experimental and molecular dynamics simulations that lipids located in one leaflet of the membrane have the capacity to interact, and in certain cases even interdigitate, with other lipids located in the opposite leaflet (i.e. the fatty acyl chains of facing membrane leaflets interact together resembling the fingers of opposing hands being locked together)^5,10–14^. This organization process, which is known as transbilayer or interleaflet coupling^15^, has been shown to impact the lateral organization of lipids in *in vitro* systems^5^. Thus, interleaflet coupling may function as an anchoring mechanism for lateral lipid partitioning, thereby potentially regulating membrane nanodomain organization and consequently cell signaling processes^5–9^.

In plants, the phytohormone auxin has been the focus of extensive research and is considered a core component of plant development and adaptation to environmental constraints. Nuclear auxin signaling triggers the transcription of auxin-responsive genes through the TRANSPORT INHIBITOR RESPONSE1 (TIR1)/AUXIN SIGNALING F□BOX (AFBs) pathway^16,17^. Apart from this transcriptional pathway, auxin activates a PM pathway through the TRANSMEMBRANE KINASE1 (TMK1) and its co-receptors to produce a rapid cell response involving protein phosphorylation and the activation of Rho-like small GTPases Rho-Of-Plants (ROP) proteins^18–24^. Upon auxin, a fraction of ROP6 proteins re-localizes in nanodomains, at the inner leaflet of the PM, that are required to trigger auxin-signaling in roots, including regulation of cytoskeleton dynamics, vesicular trafficking and PM localization of PIN2 auxin carriers^3,24^. The formation of ROP6 nanodomains in roots is subject to direct regulation by anionic lipids, with phosphatidylserine (PS) being a notable example of this regulatory mechanism^24^. It has been established that PS form constitutive nanodomains at the inner leaflet of the PM that recruit ROP6 upon auxin through a lysine/arginine motif contained in ROP6 C-terminal tail^24^. Single molecule imaging showed that about 50% of PS sensors localize in these nanodomains, where they are immobile^24^. However, how PS nanodomains are formed, and how PS molecules are blocked from diffusing in the PM remain unknown.

It is noteworthy that a distinctive feature of plant PS, in contrast to other kingdoms, is the presence of a very-long-chain fatty acid (VLCFA) with a carbon chain length of up to 24 atoms (C24)^25–27^. With the exception of PS in plants, VLCFAs are found in sphingolipids in substantial quantities, a feature shared between animals, yeasts and plants^28–34^. In both animals and plants, sphingolipids are predominantly present in the outer leaflet of the plasma membrane, while PS is located in the inner leaflet^35–43^. In accordance with the interleaflet coupling postulate, the presence of VLCFAs in the plant’s PS and sphingolipid pools has the potential to induce an orthogonal organization of the membrane. We hypothesize that this organization could stabilize PS at the PM and directly control auxin-mediated ROP6 signaling nanodomains. The potential role of this novel conceptual paradigm in auxin signaling was investigated in this study through a multidisciplinary combination of *in vivo* advanced super-resolution microscopy and computational molecular dynamics simulations.

## Results

### Molecular dynamics of interleaflet coupling through lipid interdigitation in simulations of the composition of the plant plasma membrane

Molecular dynamics simulations of animal cell PM have shown that sphingolipids in the outer leaflet can interdigitate with the opposite leaflet and interact preferentially with PS^11,12^. However, the lipid composition of plant PM differs from that of animal cells in three ways that are relevant to this study: (i) the polar head of the glycosylated sphingolipid Glycosyl-Inositol-Phosphoryl-Ceramide (GIPC) in plants contains an inositol phosphate group grafted onto a ceramide backbone, as well as the addition of a glucuronic acid and a mannose residue (**Fig. 1c**); (ii) the amidified VLCFA in plant GIPC is α-hydroxylated (**Fig. 1c**); and (iii) plant PS contains up to 50% of VLCFAs at the PM^44^. Thus, based on previously published PM lipid composition^44^, we built an all-atom model of a simplified plant PM containing 38% phosphatidylcholine (PC), 20% GIPC, 10% PS, and 32% sterols. All-atom simulations were run for 2 µs in four different combinations of acyl chain lengths: (i) the control condition containing GIPC with an α-hydroxylated fatty acid chain of 24 atoms of carbon (24;O-GIPC) and PS with a non-α-hydroxylated sn-2 fatty acid chain of 24 atoms of carbon (24-PS) (**Fig. 1a, c, Table 1, Table 2**); (ii) a condition where only the sn-2 acyl-chain of PS was shortened (**Extended data 1a, Table1, Table 2**); (iii) a condition where only the acyl-chain of GIPC was shortened (**Extended data 1b, Table1, Table 2**); and (iv) a condition where both the PS and the GIPC were shortened (**Fig. 1d, Table1, Table 2**). Since sphingolipids are located in the outer leaflet and PS in the inner leaflet^40–42,45^, we therefore placed 100% of GIPC (green, **Fig. 1a**) in the outer leaflet and 100% of PS (purple, **Fig. 1a**, **Table 1**, **Table 2**) in the inner leaflet. Sterols were distributed equally between them while PC was distributed at a ratio around 40:60 between the two leaflets, depending on the acyl chain length combination (**Table 1**, **Table 2**). Before running those simulations, we first needed to equilibrate symmetric membranes to evaluate the area per lipid of each leaflet in order to prevent that tension issues were arising between the membrane leaflets in asymmetric simulations (**Table 2**). We ran three replicates for each simulation. To test for the potential interdigitation of sn-2 24-PS and 24;O-GIPC, we quantified the density of the terminal carbons of either PS or GIPC acyl chains along the thickness of the modelled plasma membrane (0 represents the mid-plane of the plasma membrane). Our results show that the terminal carbons of PS (shown in bold purple) cross the outer leaflet, whereas the terminal carbons of GIPC (shown in bold green) cross the inner leaflet (**Fig. 1b, Extended data 1c**). Neither short-PS, short-GIPC nor 18-PC (the main phospholipid, as a control) displayed this level of interdigitation, demonstrating that the interdigitation between the two leaflets of the PM relies on VLCFAs in the PS and GIPC pools (**Fig. 1d, e, Extended data 1c**). Next, we showed that the lateral mobility of the lipids in the inner leaflet is substantially higher than those in the outer leaflet of the PM by calculating the lateral diffusion coefficient (Dlat) of lipids (**Fig. 1f**). These results are consistent with previous findings showing that outer but not inner leaflet proteins have constrained mobility^46^. Overall, the molecular dynamics results revealed an interleaflet coupling between the slow-mobile outer leaflet and the fast-mobile inner leaflet of the PM, which depends on the acyl chain length of PS and/or sphingolipids.

**Figure 1.**
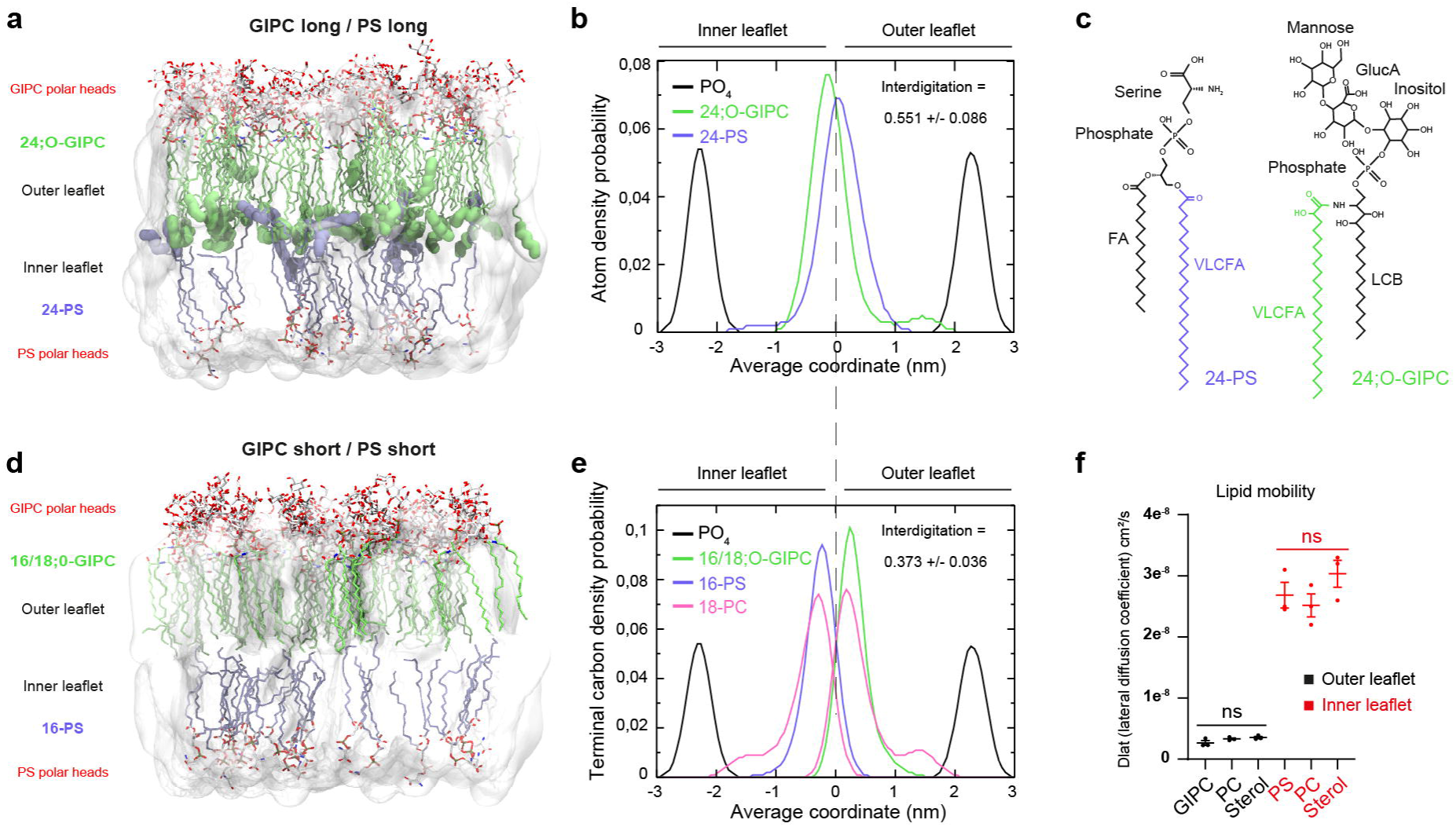
Interleaflet coupling depends on VLCFA-sphingolipids and VLCFA-PS. (**a, d**) Molecular dynamics (MD) simulations (2 µs) were performed on a lipid composition that mimics the plant plasma membrane (PM). (**a**) Control condition which contained outer leaflet-localized Very Long Chain Fatty Acid (VLCFA)-GIPC (green, 24 carbons α-hydroxylated (24;O-GIPC)) and inner leaflet-localized VLCFA-phosphatidylserine (PS) (purple, 24 carbons (24-PS)) (GIPC long / PS long). (**d**) MD simulation on a condition in which the acyl-chain of both the GIPC and PS had been reduced (GIPC short / PS short, **Table1, Table 1**). The six terminal carbons of the 24;O-GIPC and 24-PS are highlighted in bold, the polar heads are highlighted in red. (**c**) Molecular structure of 24;O-GIPC and 24-PS. While 24-PS has a short fatty acid (FA) and a non-hydroxylated VLCFA, 24;O-GIPC has a hydroxylated VLCFA and a long chain base (LCB). The polar head of PS is constituted from a phosphate and a serine while the polar head of GIPC is composed from a phosphate, inositol, glucuronic acid (GlucA) and a mannose residue. (**b, e**) Atom density probability of the presence of the terminal carbon of either the GIPC acyl-chain (green) or the PS acyl-chain (purple) according to the average coordinate across the two leaflets. It is clear that, in control condition (**b**), the terminal carbon of the 24;O-GIPC and 24-PS molecules, which are localised to the outer and inner leaflets respectively, largely overlap and cross the respective opposite leaflets. (**e**) When the acyl chains of both GIPC and PS are short, their terminal carbon only weakly interdigitate and cross opposite leaflets. In this scenario, the terminal carbon of GIPC and PS remain in a position similar to that of 18:0-PC. (**f**) The mobility of lipids in the simulations was investigated by calculating the mean square displacement of lipids (cm²/s) in the outer (black) and inner (red) leaflets. The inner leaflet is much more fluid than the outer leaflet. n=3 independent simulations. Statistics were performed by ANOVA Kruskal-Wallis, ns *P* >0.05.

**Table 1.** Description of the lipid molecular species used in the molecular dynamics simulations of Fig. 1, Fig. 5 and Extended data 1.

**Table 2.** Lipid composition used in the molecular dynamics simulations of Fig. 1, Fig. 5 and Extended data 1. The first tab provides a detailed description of the number of lipids for each lipid molecular species as well as the values of mobility in nanoseconds (ns), area in nm² and confidence interval (CI). The second tab is a synthesis of the area of lipids in simulated outer leaflet and inner leaflet of symmetric or asymmetric membranes.

### The acyl-chain length of lipids is involved in PS and ROP6 mobility at the plasma membrane

We then tested the model’s prediction of lipid interleaflet coupling *in vivo*. Having determined that interleaflet coupling depends on the length of the acyl chains of the lipids, we designed an experimental condition involving shortened acyl chains. VLCFA are synthesized by the elongase complex through a four-step enzymatic cycle involving the condensation of two acyl-CoA groups per cycle by a β-ketoacyl-CoA synthase (KCS) enzyme, followed by reduction and dehydration^47,48^. There are 21 KCS genes in Arabidopsis that display highly redundant functions, requiring multiple mutant combinations to visualize the phenotypic effects^49^. Thus, we used metazachlor (Mz), a potent and specific inhibitor of KCS enzymes, for which genetic validation had previously been provided^34,50^. Mz alters the composition of lipid pools containing VLCFAs without affecting their total quantity^34,50^. Upon Mz treatment, we previously observed that C24-VLCFAs decrease in the sphingolipid and phospholipid pools^34,50^. To confirm that this change in composition is indeed occurring in PS and GIPC at the PM, we performed lipidomic analyses of purified PM from wild-type seedlings in control condition or Mz treatment. Our results indicate that the amount of 24-PS and 24;O-GIPC decreases with Mz while the amount of 18-PS, 16;O-GIPC and 20;O-GIPC increases (**Extended data 2a, b**). Similarly, the composition profile of other anionic phospholipids besides PS, including phosphatidic acid (PA), phosphatidylinositol (PI), phosphatidylinositol phosphate (PIP) and phosphatidylinositol bisphosphate (PIP_2_), remains overall unchanged upon Mz treatment, exception made for one lipid species in the PI pool and one in the PA pool (**Extended data 2c-f**).

Having validated the specific effect of Mz on lipid chain length of PS and GIPC at the PM, we evaluated PS mobility in the PM by Fluorescence Recovery After Photobleaching (FRAP) using the PS fluorescent biosensor mCitrine-C2^LACT^ as a proxy^51^. We observed that the plateau of the recovery curve was higher in seedlings grown on Mz plates (**Fig. 2a-d**). These results suggest that the proportion of PS in the mobile fraction is higher with Mz, which supports the idea that VLCFAs play a role in anchoring PS in a slow-diffusible population. To determine whether this effect is specific to PS, we examined other anionic phospholipids that do not contain VLCFA, including phosphatidylinositol-4-phosphate (PI4P) by the use of mCitrine-1xPH^FAPP1^, mCitrine-2xPH^FAPP1^ and mCitrine-3xPH^FAPP1^ (diffusion rate similar to PS_ mCitrine-C2^LACT^, **Extended data 3a** as well as mCitrine-P4M^SidM^ (fast diffusion, **Extended data 3a**) biosensors. However, we did not observe any significant effect of Mz on the recovery curves of these PI4P biosensors (**Fig. 2e-h, Extended data 3b-e**). Furthermore, we tested two synthetic minimal protein markers that are specifically anchored at the inner leaflet of the plasma membrane (PM) through either acylation (the Myristoylated and Palmitoylated MAP-GFP) or prenylation (GFP-PAP)^46^. Both MAP and PAP display fast diffusion (**Extended data 3f**). We observed no effect of Mz on the mobility of either of these two inner-leaflet markers (**Fig. 2l, Extended data 3g, h**). REMORIN1.2 (GFP-REM1.2) and REMORIN1.3 (GFP-REM1.3), two myristoylated and palmoylated endogenous proteins arranged in nanodomains within the inner leaflet of the PM that display a slower fluorescence recovery (**Extended data 3f**)^52^ were not affected by Mz (**Fig. 2i-k, Extended data 3i, j**). Finally, we tested the mobility of three transmembrane proteins, including the syntaxin YFP-NPSN12 with one transmembrane domain (TMD), the aquaporin YFP-PIP1;4 with six TMDs, and the auxin carrier PIN2-GFP with ten TMDs that display very slow recovery curve (**Extended data 3k**). In neither case, did we observe a significant effect of Mz on the mobility of these TMD proteins (**Fig. 2m-p, Extended data 3l-o**). These results show that VLCFAs depletion affects the mobility of the anionic lipid PS in a selective manner in the inner leaflet. However, we show that a marker of the outer leaflet, the minimal GPI-anchored protein GPI-GFP^46^, displays a higher mobility upon Mz treatment suggesting that interleaflet coupling does not only impact PS in the inner leaflet but also GPI in the outer (**Extended data 4a-d**). In resting conditions, PS is partitioned between a slow-diffusible population organized into nanodomains and a more freely diffusible population evenly distributed at the PM^24^. To test if those two distinct populations of PS molecules are sensitive to acyl chain shortening, we used super-resolution single-particle tracking PhotoActivated Localization Microscopy (SPT-PALM) to track individual PS molecules within the PM^53^. We retrieved the apparent diffusion coefficient of the PS biosensor mEOS2-C2^LACT^ from mean squared displacement curves. Our results suggest that Mz treatment significantly increases the mobility of mEOS2-C2^LACT^ in the high-diffusible fraction (**Extended data 5a, b**). The SPT-PALM results corroborate the FRAP data, confirming the role of VLCFAs in restricting PS mobility at the PM. Given that ROP6 interacts with PS through its polybasic C-terminal tail^24^, we also calculated the apparent diffusion coefficient of mEos-ROP6 under control conditions (i.e. in the absence of auxin stimulation), in which ROP6 is almost exclusively present in the high-diffusible population^24^. Our results show an increase in ROP6 mobility upon Mz treatment (**Extended data 5c, d**). These results show that VLCFAs play a key role in restricting the mobility of ROP6 already in resting condition.

**Figure 2.**
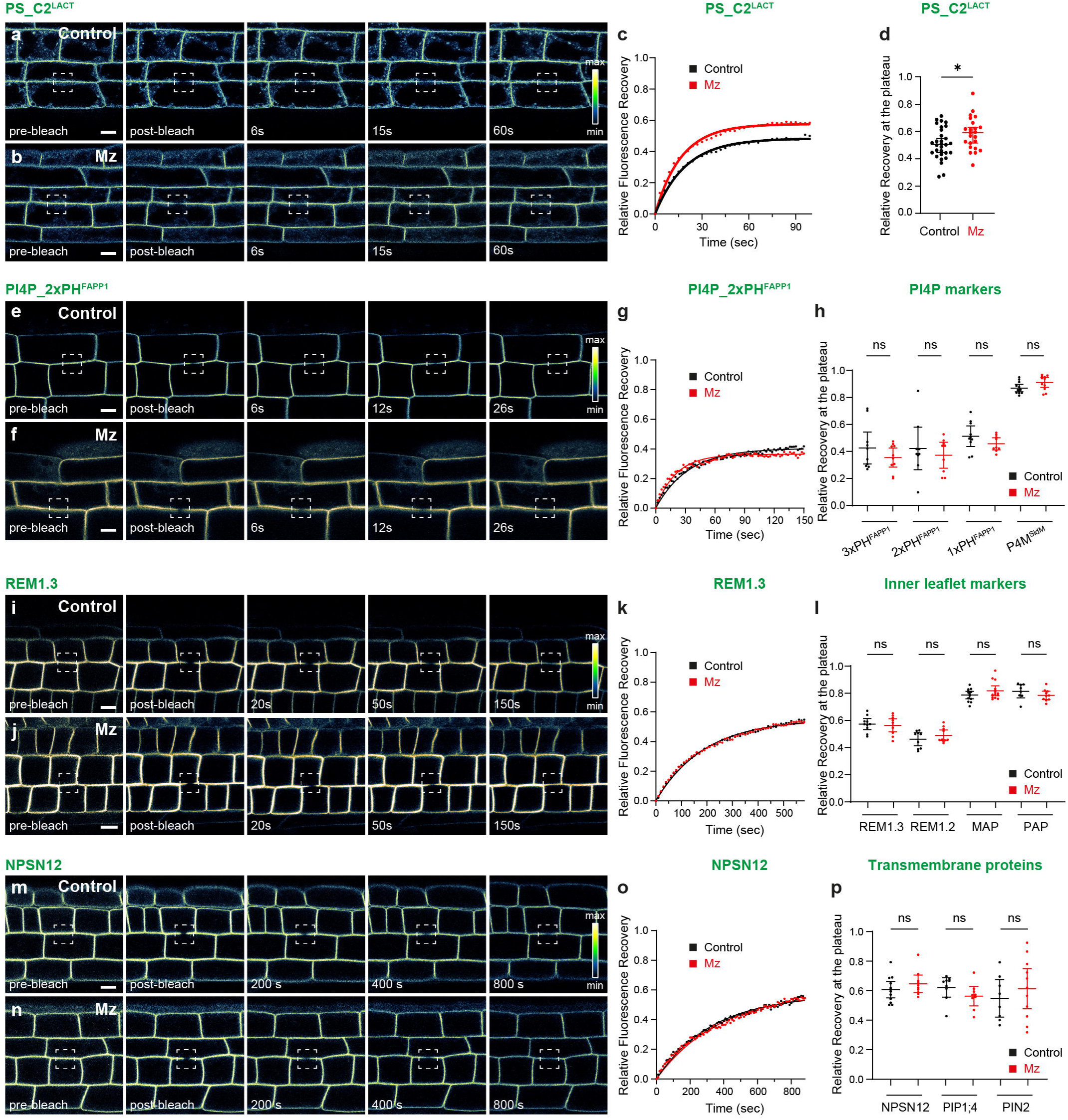
The mobility of PS in the PM is selectively dependent on VLCFA. Fluorescence Recovery After Photobleaching (FRAP) experiments in root epidermal cells of plants expressing either the inner leaflet PS biosensor mCitrine-C2^LACT^ (**a, b**), the inner leaflet PI4P biosensor mCitrine-2xPH^FAPP1^ (**e, f**), the inner leaflet YFP-REMORIN1.3 (REM1.3) protein (**i, j**) or the transmembrane syntaxin YFP-NPSN12 protein (**m, n**). After bleaching a small portion of the PM (white box in pre-bleach and post-bleach), the recovery of fluorescence was measured over time (seconds). (**a, e, i, m**) Control condition (black) *vs* (**b, f, j, n**) plants treated for 5 days on plate with the VLCFA inhibitor Metazachlor (Mz, red). (**c, g, k, o**) Relative fluorescence recovery curves corresponding to **a, b, e, f, i, j, m, n**. (**d, h, l, p**) Relative fluorescence recovery at the plateau corresponding to the curves in **c, g, k, o**. As compared to the control condition, PS (**c, d**) displayed an enhanced mobility when plants were treated with Mz (n=24-30). To know whether this effect is selective or not, (**h**) Four biosensors of PI4P were tested: mCitrine-3xPH^FAPP1^, mCitrine-2xPH^FAPP1^, mCitrine-1xPH^FAPP1^ and mCitrine-P4M^SidM^, none of them displayed significant differences between the control condition and Mz-treated plants (n=9-12). (**l**) Four protein markers of the inner leaflet were tested: the endogenous proteins YFP-REM1.3 and YFP-REM1.2 as well as the minimal Myristoylated and Palmitoylated MAP-GFP and the minimal Prenylated GFP-PAP, none of them displayed significant differences between the control condition and Mz-treated plants (n=8-14). Three endogenous transmembrane proteins were tested: the syntaxin YFP-NPSN12, the aquaporin YFP-PIP1;4 and the auxin efflux carrier PIN2-GFP, none of them displayed significant differences between the control condition and Mz-treated plants (n=8-11). Statistical tests were performed by t-test, ns *P* >0.05, * *P* <0.05, ** *P* <0.01, *** *P* <0.001, **** *P* <0.0001. All scale bars are 10 µm.

### Nano-clustering of PS and ROP6 as well as downstream auxin signaling depend on the acyl-chain length of lipids

Because VLCFA directly acts on PS diffusion in the PM, we wondered if constitutive PS nanodomains and therefore auxin-induced ROP6 nanodomains depend on lipid interleaflet coupling. Thus, we used Total Internal Reflection Fluorescence (TIRF) microscopy of root epidermal cells from seedlings expressing either the inner leaflet PS fluorescent biosensor mCitrine-C2^LACT^ or the inner leaflet GFP-ROP6 fluorescent marker. Our results clearly show that while Mz does not modify ROP6 localization pattern in resting conditions (**Fig. 3a, b, g**), it substantially decreases the density of auxin-induced (10 µM IAA for 30 min) ROP6 nanodomains (**Fig. 3d, e, g**). As Mz directly targets KCS enzymes, we examined root cells of the *kcs1* mutant, which was previously shown to downregulate VLCFA level^54^. We could confirm that auxin-induced ROP6 nanoclusters were strongly impaired in *kcs1* mutant (**Fig. 3c, f, g**). Finally, we examined PS nanoclustering using the PS fluorescent biosensor mCitrine-C2^LACT^ and TIRF microscopy. Our results show that the density of constitutive PS nanoclusters decreased significantly in the presence of Mz, or in the *kcs1* or *kcs9* mutants (**Fig. 3h-l**). Interestingly, we observed a significant increase in the density of PS nanoclusters following auxin treatment, suggesting that auxin stimulates the formation of PS nanoclusters (**Fig. 3m, n, q**). Notably, this increase in PS nanoclusters in response to auxin was prevented in both *kcs1* and *kcs9* (**Fig. 3o, p, q**). These results suggest that VLCFAs are essential for the lateral organization of PS in nanoclusters and facilitating the recruitment of ROP6 into these domains. In addition, in animal cells, PS in the inner leaflet of the PM plays a role in the nanoclustering of GPI-anchored proteins in the outer leaflet of the PM^6^. Given that we observed an increase of the lateral mobility of the outer leaflet minimal GPI-GFP marker upon Mz treatment (**Extended data 4a-d**), we then checked the localization of GPI-GFP by TIRF. Our results show that reducing the level of interleaflet coupling by Mz treatment decreases the nanoclustering of the GPI-anchored marker (**Extended data 4e-g**). These results suggest that the interleaflet coupling is not only involved in the lateral segregation of inner leaflet PS and ROP6 but also in the lateral segregation of outer leaflet GPI proteins.

**Figure 3.**
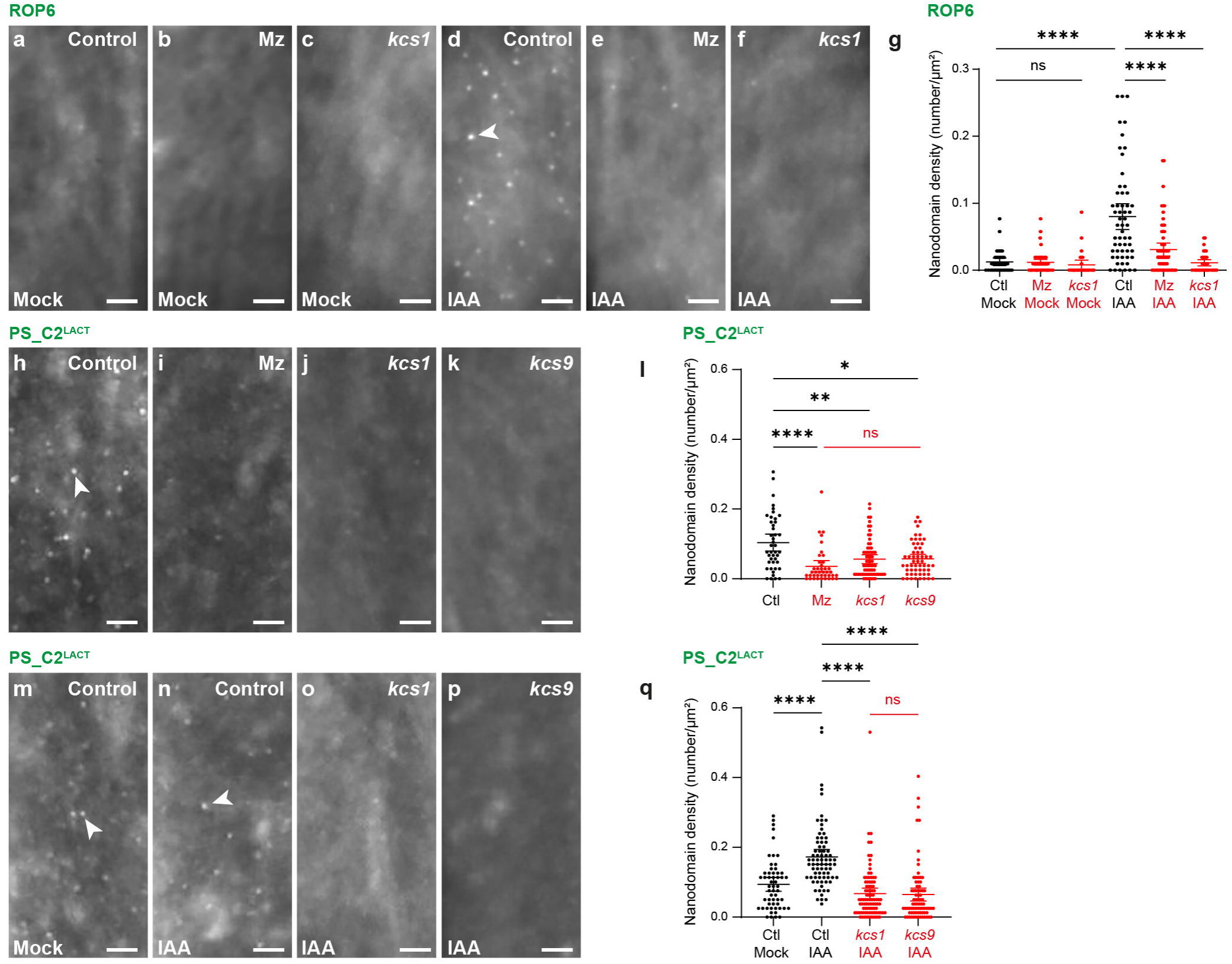
The formation of PS and auxin-induced ROP6 nanodomains is VLCFA-dependent. All images were acquired at the PM surface of root epidermal cells using Total Internal Reflection Fluorescence (TIRF) microscopy. (**a-g**) Auxin-induced ROP6-GFP nanodomains. Without auxin (mock, **a-c**), ROP6 nanodomains were not visible in the control condition (**a**), Mz treatment (**b**) or *kcs1* mutant (**c**). Contrastingly, upon auxin (IAA) stimulation (**d-f**), ROP6 nanodomains formed in the control condition (**d**, arrowheads) but not in Mz (**e**) or *kcs1* mutant (**f**), even upon auxin stimulation. (**g**) Quantification of the nanodomain density (number / µm²) corresponding to **a-f** (n=37-58). (**h-l**) PS constitutive nanodomains. In control condition (**h**), PS nanodomains are visible without auxin treatment (arrowhead in **h**) but not in Mz (**i**) or in the *kcs1* (**j**) or *kcs9* (**k**) single mutants. (**l**) Quantification of the nanodomain density (number / µm²) corresponding to **h-k** (n=40-67). (**m-q**) auxin-induced PS nanodomains visualized by the PS biosensor mCitrine-C2^LACT^. Without auxin (mock, **m**), constitutive PS nanodomains are visible (arrowhead in **m**) but their density significantly increases upon auxin stimulation (**n**, arrowheads). This effect of auxin on the density of PS nanoclusters is not observed in *kcs1* (**o**) or *kcs9* (**p**) single mutants. (**q**) Quantification of the nanodomain density (number / µm²) corresponding to **m-p** (n=55-88). Statistical tests were performed by ANOVA Kruskal-Wallis, ns *P* >0.05, * *P* <0.05, ** *P* <0.01, *** *P* <0.001, **** *P* <0.0001. All scale bars are 2 µm.

We next sought to determine whether VLCFAs could influence ROP6-mediated downstream cellular responses to auxin. One such response is the reorganization of the microtubule array^24,55,56^. Our results showed that, under control conditions, microtubules align neatly perpendicular to the growth axis (**Fig. 4a**). Treatment with Mz does not impact this orientation, as determined by calculation of the anisotropy index or orientation angle of microtubules (**Fig. 4b, e, f**). By contrast, the application of exogenous auxin (10 µM IAA for 30 min) induced the reorientation of microtubules in root cells (**Fig. 4c, e, f**). Importantly, this effect is partially prevented by Mz treatment (**Fig. 4d, e, f**). Collectively, these findings demonstrate that the presence of VLCFAs in PS and sphingolipids is essential for the formation of both constitutive PS- and auxin-induced ROP6 nanoclusters, as well as for the activation of downstream ROP6-dependent auxin signaling pathways.

**Figure 4.**
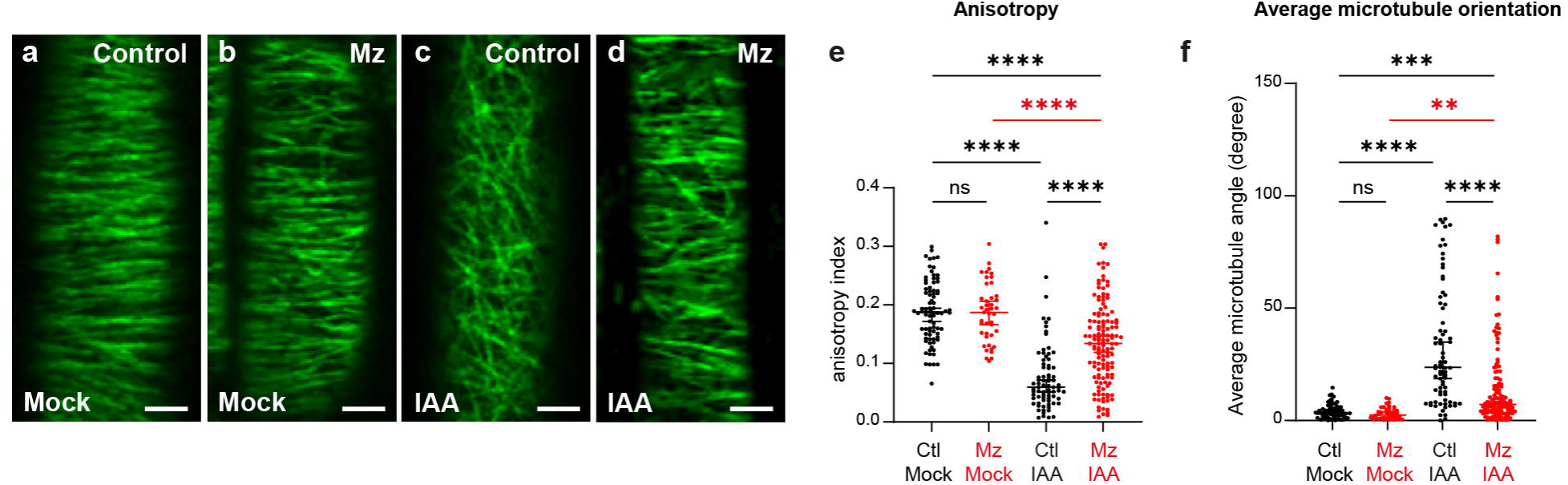
Reorientation of microtubules, downstream of auxin-signaling, is dependent on VLCFA. (**a-d**) Root epidermal cells expressing the microtubule marker TUBULIN-A6 (TUA6)-GFP in (**a**) control condition, where a nice horizontal alignment of microtubules is observed perpendicularly to the growth axis, (**b**) in Mz that does not affect microtubules organization, (**c**) upon auxin treatment, that causes a disorganization of microtubules and (**d**) upon auxin in Mz-treated plants. (**e, f**) Quantifications of the microtubule anisotropy (**e**) or average orientation angle (**f**), as compared to a horizontal reference line, corresponding to **a-d**. While auxin decreases the anisotropy of microtubules or increases the orientation angle relative to the horizontal axis, Mz partially prevents these effect. n=43-132. Statistical tests were performed by ANOVA, ns *P* >0.05, * *P* <0.05, ** *P* <0.01, *** *P* <0.001, **** *P* <0.0001. All scale bars are 2 µm.

### VLCFA-sphingolipids impact the acyl-chain ordering and mobility of PS as well as auxin-induced ROP6 nanoclustering

All-atom simulations enabled us to identify interleaflet coupling between the two leaflets of the PM, involving the terminal carbons of PS and sphingolipids (**Fig. 1a, b**). The *in vivo* experiments confirmed that VLCFAs are needed for PS lateral diffusion and nanodomain formation, thus validating our simulations. We furthered this analysis by calculating the order parameter of each carbon along the acyl chain of the different lipid species. The lipid order parameter, *S_CH_*, quantifies the average orientation and fluctuations of a given carbon–hydrogen bond in relation to the perpendicular axis of the bilayer^57^. High values indicate a more ordered lipid segment (and therefore less mobility), while low values suggest greater orientational freedom. Previous studies have shown that the order of terminal carbons in lipid chains is lower than that of carbons close to the polar head or in the middle of the chain^57,58^. Our molecular dynamics analyses show that 24-O-GIPC loosens the middle of all lipids’ acyl chains (**Fig. 5a-e**). Contrarily, in the case of interleaflet coupling, one would expect that the terminal carbons of the inner leaflet PS would be stabilized by those of the outer leaflet GIPC — or vice versa — and thus display a higher order parameter. Indeed, our *in silico* analyses revealed that the order parameter of the terminal carbons of the 24:0-PS acyl chain is higher in a membrane lipid composition containing 24;O-GIPC than in one containing short-GIPC (**Fig. 5a, b**). This effect of 24;O-GIPC on PS was not observed in a membrane lipid composition containing 18:0-PS (**Fig. 5b**). Oppositely, the order parameter of the terminal carbons of 24;O-GIPC acyl chain is not impacted by the acyl chain length of PS (**Fig. 5c)**. Importantly, neither 24;O-GIPC nor 24-PS have an impact on the order parameter of the terminal carbons of 18-PC (**Fig. 5d, e**). These results suggest that 24;O-GIPC rigidify the acyl chain of 24-PS but not the opposite way around. This could be explained by the fact that GIPC are much less mobile than PS (**Fig. 1f**). Moreover, a significant interaction was detected between GIPC and sterols in the outer leaflet (**Fig. 5f**). This specific interaction could provide an additional layer of stabilization for GIPC. Overall, these results suggest that the slow diffusion of GIPC in the outer leaflet could restrict the motion of PS terminal carbons by acyl-chain interdigitation. This would propagate along the PS acyl chain in the inner leaflet of the PM, stabilizing PS in nanodomains.

**Figure 5.**
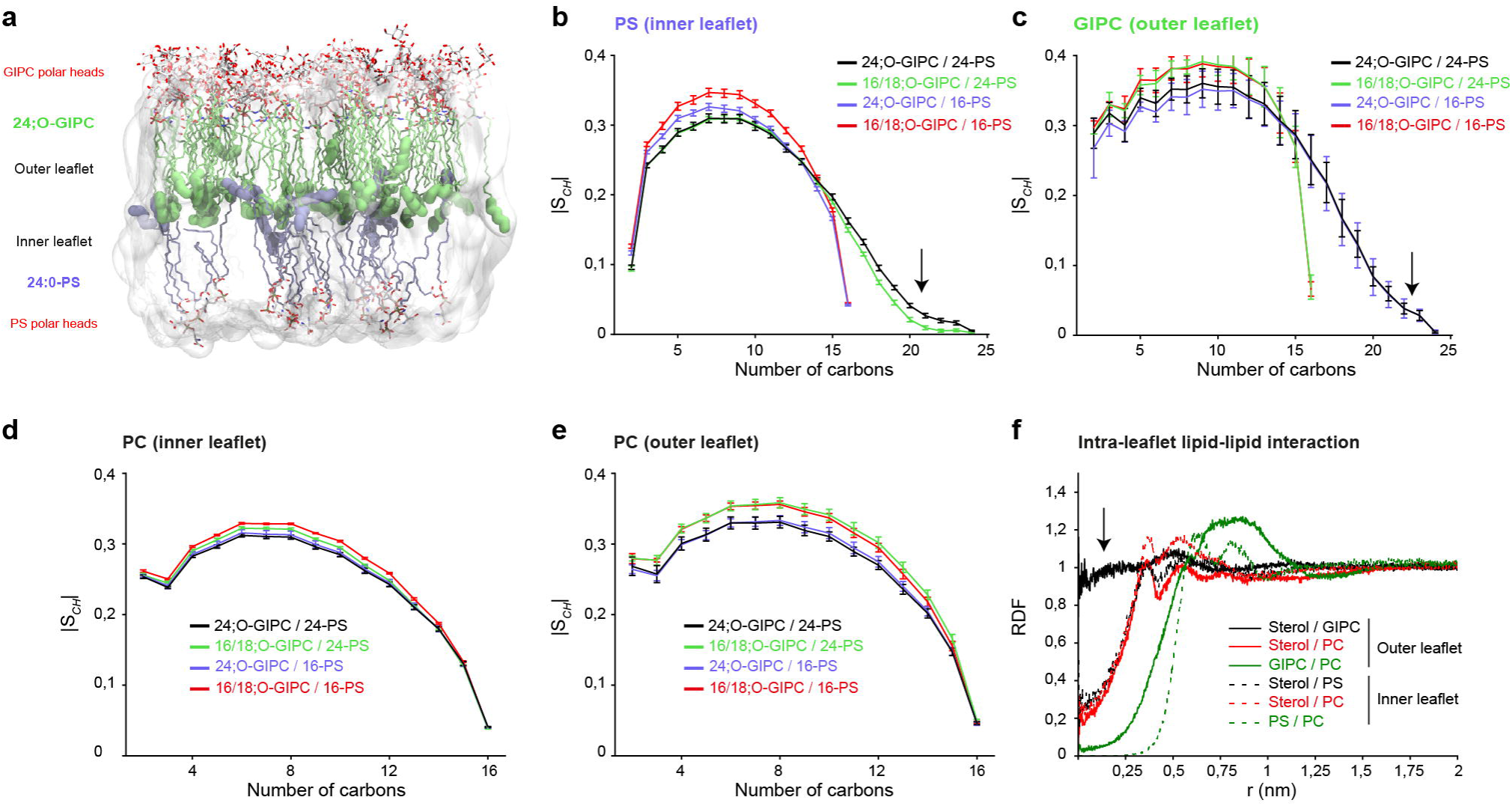
VLCFA-GIPC increases the order of the terminal carbons of VLCFA-PS. (**a**) From the same molecular dynamics simulations (2 µs) presented in Fig. 1, we calculated the order parameter (S*_CH_*) for each carbon of the acyl-chain of PS (**b**), GIPC (**c**) or PC in the inner leaflet (**d**) or outer leaflet (**e**) in either control condition (black, 24;O-GIPC / 24-PS), when GIPC acyl-chain was shortened (green), when PS acyl-chain was shortened (purple) or when both GIPC and PS acyl-chains were shortened (red). Interestingly, the order parameter of the terminal carbons of PS decreases when the acyl-chain of GIPC is shortened (**b**), meaning that the terminal carbons of GIPC positively order the terminal carbons of PS. This effect is not true opposite way around (**c**), *i.e* PS does not impact the order of GIPC terminal carbons. GIPC terminal carbons do not impact the order of PC terminal carbons (**d, e**) or PS when its acyl-chain is short (purple in **b**), suggesting some kind of selectivity of VLCFA-GIPC on VLCFA-PS. (**f**) intra-leaflet lipid-lipid interactions showing that only GIPC-sterol in the outer leaflet display a significant lipid-lipid interaction. n=3 independent simulations.

To test this hypothesis, we performed fatty acid feeding experiments. We used commercially available lignoceric (tetracosanoic) acid 24:0, which we applied directly to seedlings for 24 hours in a liquid medium in presence of Mz. To analyze whether the levels of VLCFA-PS and VLCFA-GIPC had been restored, we performed LC-MS/MS analyses on root tissues only. Our results showed that while the application of exogenous VLCFA 24:0 rescued the 24;O-GIPC pool (**Extended data 6a**), it did neither rescue the biosynthetic intermediates of GIPC (such as ceramides (**Extended data 6b**) and Inositol-Phosphoryl-Ceramides IPC (**Extended data 6c**)) or Hexose-Ceramides HexCer (**Extended data 6d**) nor did it rescue the 24:0-PS pool (**Extended data 6e**). This result may be explained by the Kennedy pathway, which uses C16/C18 fatty acid chains and CDP-DAG to synthesize phospholipids^59^. By contrast, it is well known that sphingolipids are formed by the condensation of a long-chain base (LCB) and a VLCFA^49,60^. Thus, we created an experimental condition involving VLCFA-GIPC and short-PS. In this experimental setting, we performed FRAP analyses on the mCitrine-C2^LACT^ PS biosensor. We found that, while Mz treatment significantly increased the lateral mobility of PS, exogenously applying 24:0 rescued this defect to a statistically similar level to that of untreated plants (**Fig. 6a, b, d-f**). As a control, the application of 24:0 in the absence of Mz did not affect the lateral mobility of PS (**Fig. 6c, e, f**). Additionally, we performed TIRF experiments on auxin-induced ROP6 and again found that the exogenous application of 24:0 rescued auxin-induced ROP6 nanoclustering (**Fig. 6g-l**). As a control, applying 24:0 to seedlings that had not been treated with Mz did not increase the number of auxin-induced ROP6 nanoclusters (**Fig. 6h, j, l**). Taken together, these experiments support the conclusion that the presence of VLCFA in the sphingolipid pool is necessary and sufficient to control PS diffusion and ROP6 nanodomain formation at the PM. We tested this further in the *moca1* mutant genetic background, which has a mutation in the glucuronosyltransferase involved in the grafting of the glucuronic acid residue onto inositolphosphorylceramide (IPC) to create GIPC^61^. This mutation results in a two-thirds reduction in GIPC content^61^. Our results demonstrate that the formation of auxin-induced ROP6 nanoclusters is drastically inhibited in the *moca1* mutant (**Fig. 6m-q**). These results further confirmed the role of GIPC in the formation of ROP6 nanodomains at the inner leaflet of the PM.

**Figure 6.**
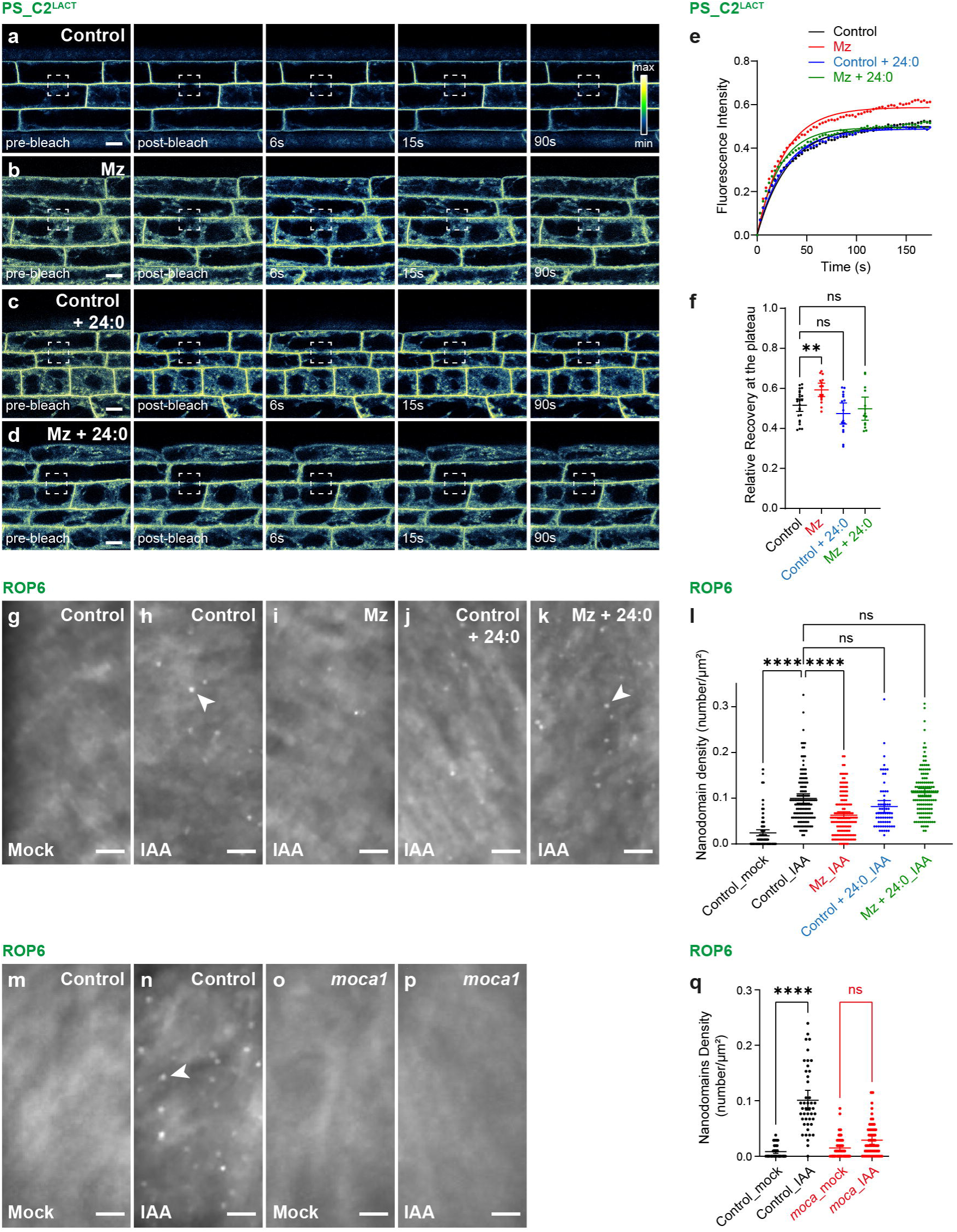
Sphingolipid-mediated PS dynamics and ROP6 nanoclustering. (**a-d**) Fluorescence Recovery After Photobleaching (FRAP) experiments in root epidermal cells of plants expressing the PS biosensor mCitrine-C2^LACT^. After bleaching a small portion of the PM (white box in pre-bleach and post-bleach), the recovery of fluorescence was measured over time (seconds). (**a**) Control condition, (**b**) plants treated for 24h with Mz, (**c**) Control condition with incubation of 24:0 fatty acid for 24h, (**d**) seedlings treated for 24h with Mz and 24:0 fatty acid. (**e**) Relative fluorescence recovery curves corresponding to **a-d**. (**f**) Relative fluorescence recovery at the plateau corresponding to the curves in **e**. As compared to the control condition (black curve), PS displayed an enhanced mobility when plants were treated with Mz (red curve), this effect was rescued by the addition of 24:0 fatty acid (green curve) (n=14-24). 24:0 fatty acid does not have an effect on their own in the control condition (blue curve). (**g-r**) ROP6-GFP nanoclustering at the PM surface of root epidermal cells observed by Total Internal Reflection Fluorescence (TIRF) microscopy in control condition (**g, m**), induced by auxin treatment (**h, n**, arrowheads), upon Mz and auxin treatment for 24h (**i**), in control condition supplemented by 24:0 fatty acids and auxin for 24h (**j**), upon Mz, auxin and 24:0 fatty acids for 24h (**k**, arrowheads) or in the GIPC *moca1* mutant without (**o**) or with auxin (**p**) induction. (**l, q**) Quantification of the nanodomain density (number / µm²) corresponding to **g-k** and **m-p**, respectively (n=66-145 in **l**, n=44-64 in **q**). Auxin (IAA) induces the formation of ROP6 nanodomains, a Mz treatment of 24h or the *moca1* mutant partially inhibits this process while an incubation with 24:0 fatty acid for 24h together with Mz rescue the ROP6 defect caused by Mz. Statistical tests were performed by ANOVA in **f** and by ANOVA Kruskal-Wallis in **l, q**, ns *P* >0.05, * *P* <0.05, ** *P* <0.01, *** *P* <0.001, **** *P* <0.0001. Scale bars are 10 µm in **a-d** and 2 µm in **g-p**.

## Discussion

Our understanding of the organization of biological membranes has evolved significantly since the fluid mosaic model^62^. It has become increasingly clear that the PM is not homogeneous in composition. Indeed, it has been shown that the two leaflets of the PM have an asymmetric lipid composition^35,63^. In animal cells, PS and PE are almost exclusively located in the inner leaflet, and sphingomyelin and glycosylated sphingolipids are almost exclusively located in the outer leaflet^35–39^. In plants, the lipid asymmetry between the two leaflets of the PM has been studied in mung beans by producing right-side-out and inside-out vesicles from purified PM fractions, and by analyzing the rate at which phospholipids are hydrolyzed when exposed to externally applied phospholipases^40^. These experiments revealed that while PC, PE and PA were homogeneously distributed between the two leaflets, PS was distributed asymmetrically within the inner leaflet of the PM^40,41^. Aside from phospholipids, glycosylated sphingolipids are expected to be located in the outer leaflet of the PM because the glycosylation of ceramides to produce glucosylceramide (ClcCer), or the addition of an inositolphosphate group to ceramides and further glycosylation to produce GIPC, occur in the luminal leaflet of the ER and the Golgi complex, respectively^50,64–66^. Indeed, it was observed that the GlcCer is primarily located in the outer leaflet of PM vesicles^42^. The polar head of GIPC is much larger than that of GlcCer; therefore, it is unlikely that GIPC would flip-flop between the two leaflets of the Golgi or the PM. Immuno-electron microscopy analysis of tobacco PM vesicles has consistently showed that GIPCs reside in the outer leaflet of these vesicles^43^. What is the impact of this lipid asymmetry in membrane function? Molecular dynamics simulations of the composition of animal cell membranes have suggested that there is a preference for C24-sphingomyelin in the outer leaflet to interact with PS in the inner leaflet^11^. Our molecular dynamics simulation revealed that 24;O-GIPC increase the order of the terminal carbons of PS preferentially. Additionally, using a set of PM-localized markers, we demonstrate that VLCFAs selectively alter PS mobility. Our fatty acid complementation assay and genetic approach show that 24;O-GIPC are involved in mobility and nanoclustering of PS at the PM, and thereby the formation of auxin-induced ROP6 nanodomains. Our results show that the presence of VLCFAs in PS is not necessary for this effect, but rather depends on the presence of VLCFAs in GIPC. The reason for VLCFA sphingolipids preferring PS is unclear, but this phenomenon is also observed in animal cells where it has been proposed that sphingolipids and 18-PS act together on localization of the small GTPase KRAS^67^. Separately from sphingolipids, in animal cells 18-PS plays a role in the nanoclustering of GPI-anchored proteins on the opposite side of the PM^6^. One explanation for this specificity is the interaction between PS and the actin cytoskeleton^68^. This interaction has been proposed to restrict the diffusion of PS^69^ and lead to the “picket-fence” model that is now encompassed by the “active actin-membrane composite” model, a new paradigm in which the orthogonal lipid membrane asymmetry plays a crucial role in regulating lateral nanodomains^6,70^. This model involves the asymmetric lipid composition in membrane leaflets, as well as lipid-lipid interactions across leaflets, which induce interleaflet coupling^6,70,71^. However, the impact of this on cell signaling or other mechanisms that use the PM as an integrative interface between the cell and its environment — such as cell-to-cell communication, transport, and host-pathogen interaction — has scarcely been investigated^72^. Here, we present *in vivo* and *in silico* evidence supporting the role of outer leaflet sphingolipids in mediating the formation of ordered PS nanoclusters in the inner leaflet of the PM through interleaflet coupling. Interestingly, we found that interleaflet coupling is required for forming Rho-GTPase ROP6 nanoclusters that are crucial for the auxin response and downstream signaling events, such as microtubule reorientation that support cell growth. We do not believe that interleaflet coupling is limited to organizing the inner leaflet. In fact, in animal cells, PS in the inner leaflet of the PM plays a role in the nanoclustering of GPI-anchored proteins in the outer leaflet of the PM^6^. Our results consistently show that reducing the level of interleaflet coupling decreases the nanoclustering of a minimal GPI-anchored marker. Overall, our study provides a rare insight into the role of lipid interleaflet coupling in cell signaling and suggests that similar mechanisms operate in animals and plants.

## Online methods

### Molecular Dynamics

To study the effects of lipid interdigitation on membrane properties, asymmetric membrane models containing 200 lipids have been analyzed using molecular dynamics (MD). These models contained GIPC and/or PS lipids with either a long (C24) or short (C16/C18) acyl chain. GIPC are found in the outer leaflet, while PS are found in the inner leaflet, leading to the creation of four different models. In the asymmetric membrane models, the number of lipids in each leaflet was calculated so that the surface area would be the same as that of the symmetric membranes. The lipid composition had to be adjusted depending on the acyl chain length, with three slightly different compositions being used. The lipid descriptions are given in Table 1. The compositions and simulations performed are listed in Table 2. All simulations have been performed with the CHARMM36 force field^73^. Membrane systems have been generated by using the CHARMM-GUI membrane builder^74,75^ and the box filled with TIP3P water^76^. When the lipids were not available with CHARMM-GUI, the closest matching composition was built and the lipid structures and topologies were adapted from existing ones. All the systems studied were equilibrated by using the six steps equilibration proposed by the CHARMM-GUI membrane builder; a minimization by steepest descent of 1,000 steps, two NVT and four NPT simulations with increasing length and time step and decreasing restraints force constants on lipids phosphate positions and dihedrals. Temperature and pressure were coupled at 303.15 K and 1 bar using the weak coupling Berendsen algorithm with τT = 1 ps and τP = 5 ps^77^. Pressure was coupled semi-isotropically. The production simulations were then carried out for 2000 ns. Periodic boundary conditions (PBC) are used with a 2 fs time step. Temperature was maintained by using the Nosé-Hoover method^78,79^ and pressure by using the Parrinello-Rahman barostat^80^ with a compressibility of 4.5 × 10^5 (1/bar). Electrostatic interactions were treated by using the particle mesh Ewald (PME) method^81^. The van der Waals interaction was switched off from 1 to 1.2 nm by the force-based switching function^82^. Hydrogen bonds lengths were maintained with the LINCS algorithm^83^. Trajectories were performed and analyzed with GROMACS 2020 tools^84^, MDAnalysis^85^ and LipidDyn^86^. Interdigitation is evaluated through the overlap parameter as described^11^. 3D structures were analyzed with both PYMOL^87^ and VMD^88^ software. Radial distribution functions have been computed for the last 400 ns of the trajectories with the phosphate, ceramide first carbon and sitosterol oxygen.

### Plant material and growth conditions

The following Arabidopsis transgenic fluorescent marker lines were used: pUBQ10::mCITRINE-C2^LACT^ ^51^, pUBQ10::mCITRINE-1xPH^FAPP1^ (P5Y)^89^, pUBQ10::mCITRINE-2xPH^FAPP1^ (P21Y)^89^, pUBQ10::mCITRINE-3xPH^FAPP1^ ^89^, pUBQ10::mCITRINE-P4M^SidM^ ^51^, p35S::GFP-ROP6^3^, p35s::YFP-REM1.3^52^, p35s::YFP-REM1.2^52^, p35s::MAP-GFP^90^, p35s::GFP-PAP^90^, p35s::GFP-GPI^90^, pUBQ10::YFP-NPSN12^91^, pUBQ10::YFP-PIP1;4^91^, pPIN2::PIN2-GFP^92^ and p35S::GFP-TUA6^93^. The following Arabidopsis mutants were used: *kcs1-5* (SALK_200839)^94^, *kcs9* (SALK_028563)^95^ and *moca1*^61^. For all observations, seeds were treated with 0.1% Triton in water during 5 minutes, then washed three times with water and transferred to 4°C for 2 days. The seeds were then sterilized by a 0.9% chlorine/0.1% Triton solution during 20 minutes and sown on ½ Murashige and Skoogs (MS) agar medium plates (1% plant agar, 1% sucrose, and 2.5mM morpholinoethanesulfonic acid pH5.8 with KOH). The seedlings were grown 5 or 6 days vertically in long day conditions (150 mE/m^2^/s -1, 16h light/ 8h dark) at 22°C.

### Confocal microscopy

Fluorescence Recovery After Photobleaching (FRAP) acquisitions were done using the confocal microscopy of a Zeiss LSM 880 using 40X oil-immersion objective. According to each marker dynamics, images were acquired every 3 sec up to 180 sec (60 images per experiment) for pUBQ10::mCITRINE-C2^LACT^, p35s::MAP-GFP and p35s::GFP-PAP, every 2 sec until 200 sec (100 images per experiment) for pUBQ10::mCITRINE-1xPH^FAPP1^ (P5Y), pUBQ10::mCITRINE-2xPH^FAPP1^ (P21Y), pUBQ10::mCITRINE-3xPH^FAPP1^ and pUBQ10::mCITRINE-P4M^SidM^, every 10 sec until 600 sec (60 images per experiment) for p35s::YFP-REM1.3 and p35s::YFP-REM1.2, every 10 sec until 900 sec (90 images per experiment) for pUBQ10::YFP-NPSN12 and pUBQ10::YFP-PIP1;4 and every 15 seconds until 22.5 min (90 images per experiment) for p35s::GFP-GPI. For pPIN2::PIN2-GFP, images were taken every 30 seconds until 5 min and then one image was taken every 5 min until 30 min (15 images per experiment). In all cases, seedlings were mounted between an Epredia™ SuperFrost Plus slide and a 20x20 mm #1.5 coverslip spaced by double-sided in ½ MS liquid medium (1% sucrose, and 2.5mM morpholinoethanesulfonic acid pH5.8 with KOH). The 488 and 514 nm excitation laser was used for GFP and mCitrine, respectively. Fluorescence recovery was subsequently analyzed in the bleached ROIs and in controlled ROIs (3 rectangles in unbleached area). Fluorescence intensity of an ROI was systematically measured outside the root and subtracted to plasma membrane as background correction. Fluorescence intensity data were normalized using the equation: I_n_=(I_t_-I_min_)/(I_max_-I_min_). Where I_n_ is the normalized intensity, I_t_ is the intensity at any time t, I_min_ is the minimum post-photobleaching intensity and I_max_ is the mean pre-photobleaching intensity. Normalized recovery data were then fitted to an exponential recovery fitting curve using One-Phase Nonlinear Regression in GraphPad Prism 10 (GraphPad Software, San Diego, CA, USA).

### Microtubules orientation

Microtubule array confocal images were acquired on 6-day-old seedlings in the elongation zone of root cells. The average orientation and anisotropic index were both calculated on 91, 43, 76 and 132 root cells for control, metazachlor treatment, auxin treatment and metazachlor/auxin combined treatment of 6 biological repeats using FibrilTool^96^ plugin on Fiji^97^.

### Total internal reflection fluorescence

Total internal reflection fluorescence (TIRF) was done using a custom set up made with an inverted Zeiss microscope with an oil immersion objective 100x/1.45 for a final 150-fold imaging magnification (corresponding to a pixel size of 102 nm) and equipped with an EMCCD camera (iXon X3 DU-897, Andor Technologies). For GFP imaging, excitation was set at 488 nm using a 488 nm (OBIS, LX 488-50, Coherent Inc.) laser set at 40 mW and used at 5%. Light was spectrally filtered at 525 +/- 22.5 nm using emission filter. For each acquisition, one hundred images were recorded with a 0.05 second exposure time. A Z-stack by average intensity was then realized prior detection of the clusters using a machine learning-based segmentation with ilastik software^98^.

### sptPALM

Imaging was performed on a Zeiss Elyra PS1 system with a 100x Apo (numerical aperture 1.46 Oil objective), in TIRF mode equipped with EMCCD iXon897 Ultra camera. The optimum critical angle was determined as giving the best signal-to-noise ratio. Pixel size was 0.097 μm. mEOS was photoconverted using 405 nm UV laser power and resulting photoconverted fluorophores were excited using 561 nm laser. UV laser power was adjusted to have significant number of tracks without too high density to facilitate further analyses (0.01 to 0.08%). 10000 images time series were recorded at 50 frames per second (20ms exposure time) on a 256 x 256-pixel region of interest.

### Inhibitor treatments

Metazachlor (Mz, Merck, Sigma PESTANAL®) treatment was performed on seedlings grown for 6 days on 100 µM Mz-containing ½ MS plates or 5 days on ½ MS plates and then transferred to a liquid ½ MS medium containing 100 µM Mz for 24 h. Indole-3-acetic acid (IAA; CAS No. 87-51-4, Duchefa Biochemie, Haarlem, The Netherlands) in DMSO was added to liquid media at 10 µM final concentration for 30 minutes. DMSO was used as control.

### Fatty acid add-back

For the treatment, media containing fatty acids were prepared by adding 24:0 (lignoceric acid: tetracosanoic acid, Matreya) to liquid ½ MS liquid medium and heating at 70 °C for 30 min. The fatty acids were added from 25 mM stock solutions inCHCl_3_/MeOH (5:1) solvent mix. After heating, the media were cooled to the room temperature and 100 nM Mz was added. Arabidopsis seedlings were grown on half MS plates without Mz and transferred to the liquid media containing fatty acid and Mz. They were incubated for 24 h with mild shaking under 16 h light/8 h darkness at 22 °C. The plants were directly used for the confocal or TIRF imaging after treatment. For the fatty acid quantification, the seedlings were washed by 30 mL of half MS for three times after treatment in order to remove the remaining fatty acid on the plant surface, and only the roots were collected.

### Lipid extraction and derivatization procedure

For anionic lipid extraction, we followed the protocol described previously^27^. 725 µl of a MeOH/CHCl_3_/1M HCl (2:1:0.1, v/v/v) solution and 150 µL water were added to the samples in Eppendorf tubes which were then grinded three times for 30 sec using a TissueLyser II (Qiagen, Courtaboeuf, France) with metal beads. Following the addition of the internal standard (15:0/18:1 PA, 17:0/14:1 PI, 17:0/14:1 PS, 17:0/20:4 PI4P and 17:0/20:4 PI(4,5)P2) and of 750 µl CHCl_3_ and 170 µl HCl 2 M, the samples were vortexed and centrifuged (1500 g / 5 min). The lower phase was washed with 708 µl of the upper phase of a mix of MeOH/CHCl_3_/0.01 M HCl (1:2:0.75, v/v/v). Samples were vortexed and centrifuged (1500 g / 3 min) and then washed again. Then, samples were kept overnight at 20°C. The organic phase was transferred to a new Eppendorf tube and the methylation reaction was carried out with 50 µl of TMS-diazomethane (2 M in hexane). After 10 min of reaction at 23°C, 6 µl of glacial acetic acid was added. 700 µl of the upper phase of a mix of MeOH/CHCl_3_/H_2_O (1:2:0.75, v/v/v) was added, vortexed, centrifuged (1500 g/ 3 min) and upper phase was removed. 700 µl of the upper phase of a mix of MeOH/CHCl_3_/H_2_O was added again, vortexed, centrifuged upper phase was removed. Finally, the lower organic phases were transferred to a new Eppendorf tube. Following the addition of 100 µl MeOH/H_2_O (9:1, v/v), the samples were concentrated under a gentle flow of air. 80 µl of methanol were then added, submitted to ultrasounds for 1 min. Then, 20 µl water were added and submitted to 1 more min of sonication.

Sphingolipids were extracted as described previously^44^ in the lower phase of IPA/Hexane/H_2_O (55:20:25, v/v/v) at 60°C for 1 h. The extracts were dried and resuspended in CHCl_3_/MeOH/H_2_O (30:60:10, v/v/v). For LC–MS/MS analysis, sphingolipid extracts were then incubated 1 h at 50°C in 2 ml of methylamine solution (7 ml methylamine, 33% (w/v) in EtOH combined with 3 ml of methylamine 40% (w/v) in H_2_O in order to remove phospholipids. After incubation, methylamine solutions were dried at 40°C under a stream of air. Finally, they were resuspended into 100 µL of THF/MeOH/H_2_O (40:20:40, v/v/v), 0.1% HCOOH containing synthetic internal lipid standards (Cer d18:1/h17:0, Cer d18:1/C17:0, GlcCer d18:1/C12:0) was added, thoroughly vortexed, incubated at 60°C for 20 min, sonicated 2 min, and transferred into LC vials.

### HPLC-MS/MS analysis

Analysis of methylated anionic phospholipids was performed using a high-performance liquid chromatography system (1290 Infinity II, Agilent, Santa Clara, CA, USA) coupled to a QTRAP 6500 mass spectrometer (Sciex, Framingham, MS, USA). The chromatographic separation of anionic phospholipids was performed on a reverse-phase C18 column (SUPELCOSIL ABZ+; 10 cm x 2.1 mm, 3 lm, Merck, Darmstadt, Germany) using MeOH/H_2_O (3:2, v/v) as solvent A and IPA/MeOH (4:1, v/v) as solvent B at a flow rate of 0.2 ml/min. All solvents were supplemented with 0.1% HCOOH and 5 mM ammonium formate. 20 microliters of samples were injected and the percentage of solvent B during the gradient elution was as following: 0–20 min, 45%; 40 min, 60%; 50 min, 80%. The column temperature was kept at 40°C. Analyses were performed in the positive mode. Nitrogen was used for the curtain gas (set to 35), gas 1 (set to 40), and gas 2 (set to 40). Needle voltage was at +5500 V with needle heating at 350°C; the declustering potential (DP) was +10 V. The collision gas was also nitrogen; collision energy (CE) was set between 26 and 45 eV according to the lipid classes. For quantification, the areas of LC peaks were determined using MultiQuant software (version 3.0, Sciex).

For sphingolipids, LC–MS/MS (multiple reaction monitoring mode) analyses were performed with a QTRAP 6500 (ABSciex) mass spectrometer coupled to a liquid chromatography system (1290 Infinity II, Agilent, Santa Clara, CA, USA). Analyses were performed in the positive mode. Nitrogen was used for the curtain gas (set to 30), gas 1 (set to 30), and gas 2 (set to 10). Needle voltage was at +5500 V with needle heating at 400°C; the de-clustering potential was adjusted between +10 and +40 V. The collision gas was also nitrogen; collision energy varied from +15 to +60 eV on a compound-dependent basis. Reverse-phase separations were performed at 40°C on a SUPERCOSIL ABZ+, 10 cm x 2.1 mm column and 3 µm particles (Merck, Darmstadt, Germany). The mobile phase consisted of a gradient of solvents A: THF/ACN/5 mM ammonium formate (3:2:5, v/v/v) with 0.1% HCOOH and B: THF/ACN/5 mM ammonium formate (7:2:1, v/v/v) with 0.1% HCOOH. The gradient elution program for Cer and GlcCer quantification was as follows: 0 to 1 min, 1% B; 40 min, 80% B; and 40 to 42, 80% B. The gradient elution program for GIPC quantification was as follows: 0 to 1 min, 15% B; 31 min, 45% B; 47.5 min, 70% B; and 47.5 to 49, 70% B. The flow rate was set at 0.2 ml/min, and 20 ml sample volumes were injected. For quantification, the areas of LC peaks were determined using MultiQuant software (version 3.0; Sciex).

### Statistical information

To compare three groups or more, the One-Way ANOVA test was performed if all the datasets follow a Gaussian distribution, otherwise the Kruskal-Wallis test was used. In FRAP and spt-PALM experiments, t-test was used to compare Mz effects and control condition for all markers. For lipidomics, ANOVA and Shapiro-Wilk test were performed because the number of replicates were between 3 and 6. All statistical analysis was done in GraphPad Prism 10 (GraphPad Software, San Diego, CA, USA).

## Acknowledgments

We thank Elia Stahl for critical reading of the manuscript. Imaging was performed at the Bordeaux Imaging Center (BIC), of the Montpellier Ressources Imagerie (MRI), the Histocytology and Plant Cell Imaging Platform (PHIV) and MARS. All imaging platforms are part of the National Infrastructure France-BioImaging (FBI) supported by the French National Research Agency (ANR-10-INSB-04). Lipidomics was performed at the lipidomic facility in Bordeaux, part of the Bordeaux metabolome platform and the MetaboHub national infrastructure funded by ANR (ANR-11-INBS-0010). Work in Y.B. lab was funded by the ANR FATROOT (ANR-21-CE13-0019) and the ANR PLAYMOBIL (ANR-19-CE20-0016). The authors thank the ROMEO regional calculation center at the University of Reims Champagne-Ardennes for providing computational resources and support. Work in Y.J. lab was supported by the European Research Council (ERC) under the European Union’s Horizon 2020 research and innovation program (projects 101001097-LIPIDEV). Work in A.M. lab is supported by the Agence National de la Recherche (ANR) CellOsmo (ANR-19-CE20-0008) and Nano-ROS (ANR-24-CE92-0024).

## Author contribution statement

M.M performed FRAP, TIRF and lipidomic experiments and analyses in Fig. 2, 3, 4, 6 and Extended data 2, 3, 5. A.P performed TIRF experiments and analyses in Fig. 3 and 6g-u. C.D setup the initial molecular dynamics simulations of Fig. 1, 5 and Extended data 1. V.B performed the SPT-PALM experiments in Extended data 4. M.N, A.M.C, M.D.J performed FRAP experiments in Fig. 2 and Extended data 3. J.B.F participated to TIRF experiments in Fig. 3 and 6. C.S participated in the initial molecular dynamics simulations. J.M.C designed and performed all experiments related to molecular dynamics in Fig. 1, 5 and Extended data 1, and supervised C.D. L.F run all lipid extracts on the LC-MS/MS machine, analyzed and interpreted the data. F.S.P, Y.J and S.M provided scientific input, genetic materials and technical resources essential to this study. Y.B and A.M conceptualized and designed the research, got the financial support for, supervised all aspects of this study and wrote the manuscript. The figures were made by M.M and Y.B. All authors read and provided inputs on the manuscript.

## Data availability statement

Data supporting the findings of this work are available in this paper and its extended data and supplementary information. All datasets and plant genetic materials generated and analysed in this study are available from the corresponding author upon request. Source data are provided with this paper.

## Extended data figure legends

**Extended data 1.**
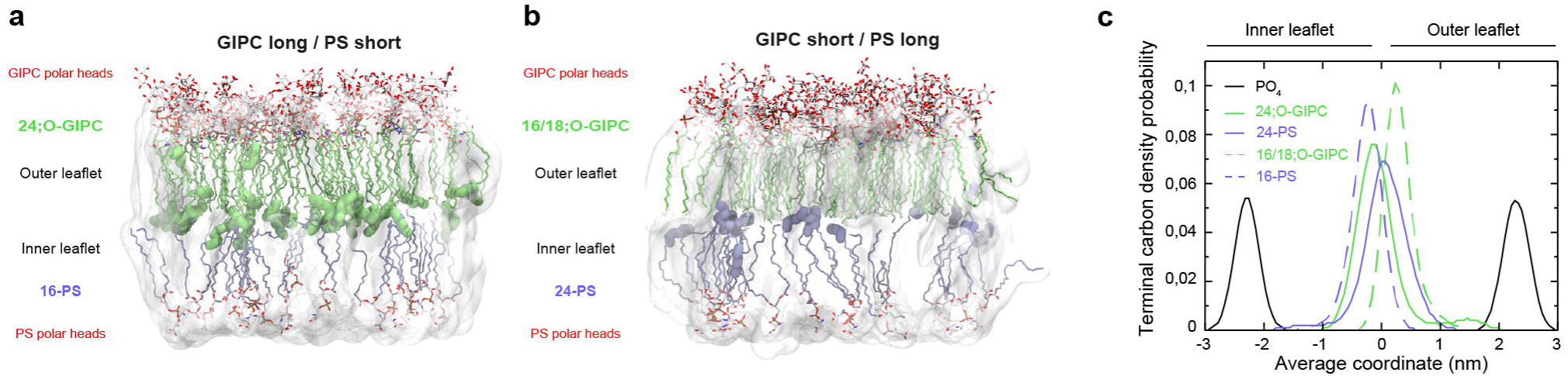
Additional results supporting Fig. 1. Interleaflet coupling depends on VLCFA-sphingolipids and VLCFA-PS. (**a, b**) Complementary molecular dynamics simulation models (2 µs) of Fig. 1a, b. (**a**) Simulation model with outer leaflet-localized Very Long Chain Fatty Acid (VLCFA)-GIPC (green, 24 carbons α-hydroxylated (24;O-GIPC)) and inner leaflet-localized short-PS (purple) (GIPC long / PS short). (**b**) Model with short-GIPC and 24-PS (GIPC short / PS long). The six terminal carbons of the 24;O-GIPC (in **a**) or 24-PS (in **b**) are highlighted in bold, the polar heads are highlighted in red. (**c**) Atom density probability of the presence of the four terminal carbons of either the GIPC acyl-chain (green) or the PS acyl-chain (purple) according to the average coordinate across the two leaflets, in four different scenarios: GIPC long, GIPC short, PS long and PS short. High degree of interdigitation is observed only when both GIPC and PS are long. n=3 independent simulations.

**Extended data 2.**
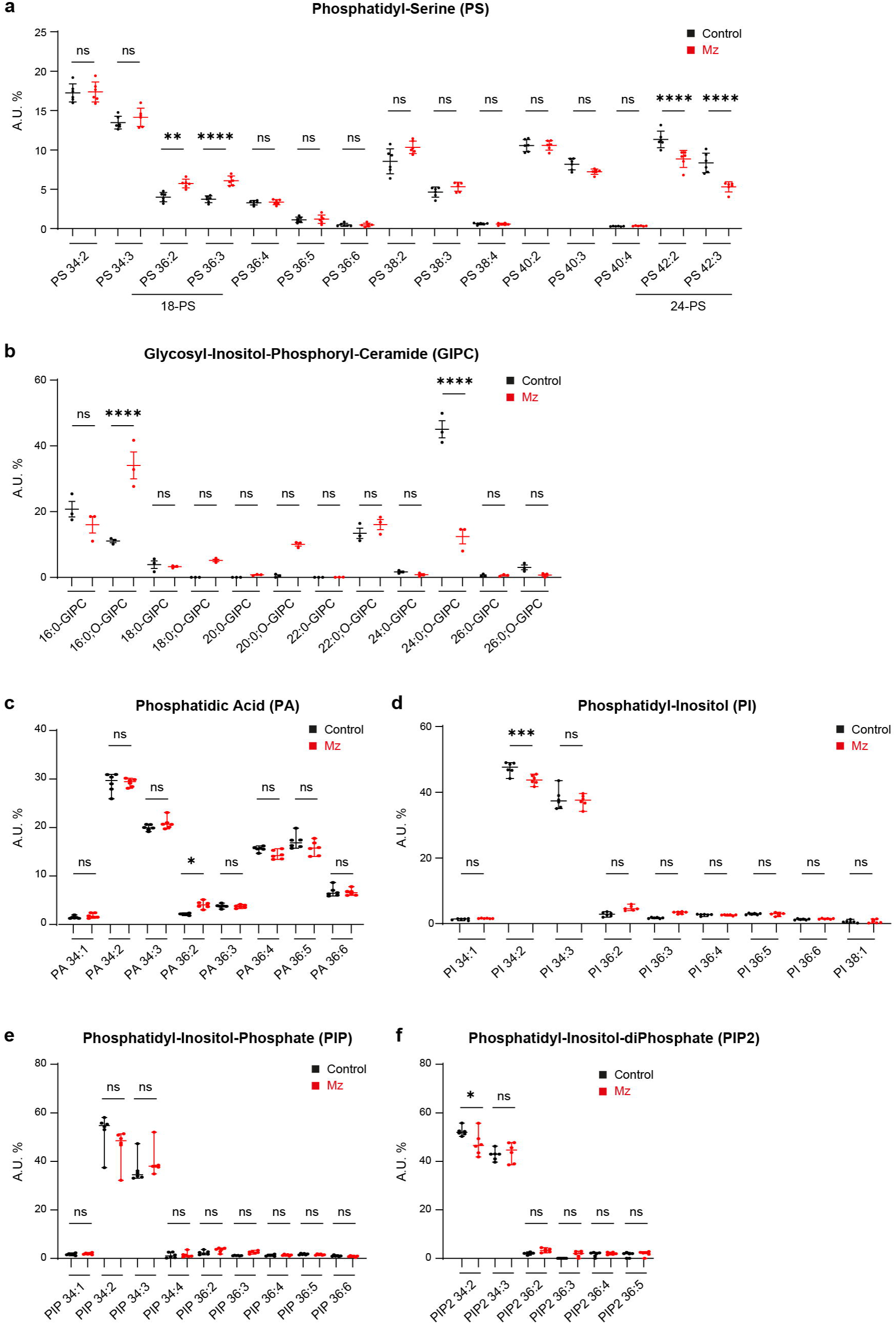
Lipidomic analysis of the plasma membrane from whole seedlings grown on Metazachlor (Mz). LC-MS/MS analyses of (**a**) phosphatidylserine (PS), (**b**) glycosyl-inositol-phosphoryl-ceramides (GIPC), (**c**) phosphatidic acid (PA), (**d**) phosphatidylinositol (PI), (**e**) phosphatidylinositol phosphate (PIP) and (**f**) phosphatidylinositol bisphosphate (PIP_2_). Upon Mz (red), the relative amount of 24-PS and 24-GIPC decreases while 16-GIPC increases (n=3). No or minor modifications were observed in the pools of PA, PI, PIP and PIP2 (n=6). Statistical tests were performed by ANOVA, ns *P* >0.05, * *P* <0.05, ** *P* <0.01, *** *P* <0.001, **** *P* <0.0001.

**Extended data 3.**
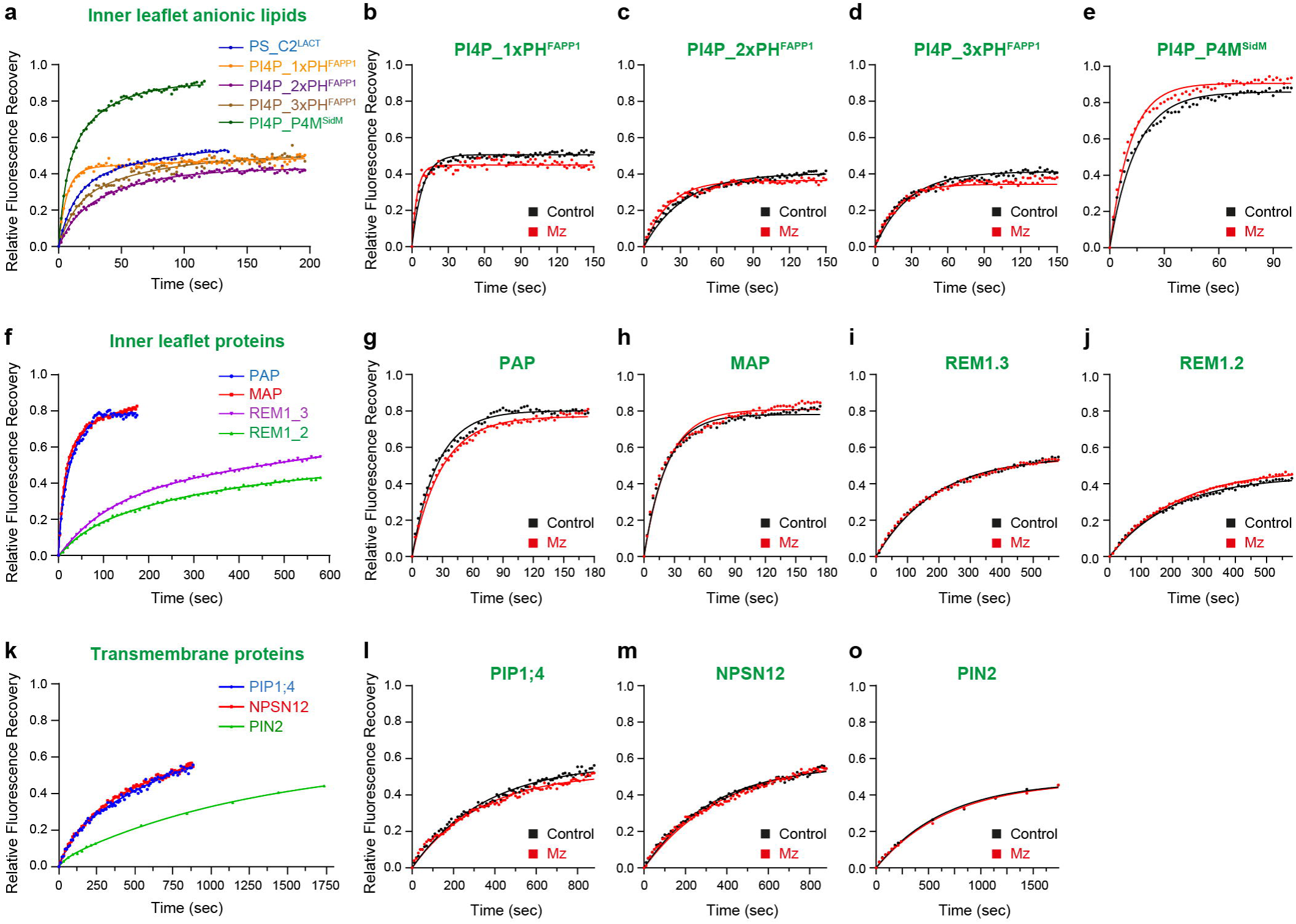
Supplementary FRAP results supporting Fig. 2. Recovery curves for all fluorescent makers used and quantified in Fig. 2h, l, p. (**a**) Comparison of fluorescence recovery for the PS biosensor C2^LACT^ and different PI4P biosensors (1xPH^FAPP1^, 2xPH^FAPP1^, 3xPH^FAPP1^, P4M^SidM^). The PI4P biosensor P4M^SidM^ displays a fast mobility at the PM while the PS C2^LACT^ biosensor and the PH^FAPP1^ biosensors show similar kinetics. (**f**) Comparison of fluorescence recovery for a set of inner leaflet proteins. The minimal Myristoylated and Palmitoylated GFP-MAP and the Prenylated PAP-GFP display fast kinetics while the endogenous YFP-REM1.3 and YFP-REM1.2 show slower kinetics. (**k**) Comparison of fluorescence recovery for a set of transmembrane proteins. As compared to the syntaxin YFP-NPSN12 and the aquaporin YFP-PIP1;4, the auxin efflux carrier PIN2 display very slow kinetics. (**b-e, g-j, l-o**) Relative fluorescence recovery curves of different markers in control condition plants (black) *vs* plants treated for 5 days on plate with the VLCFA inhibitor Metazachlor (Mz, red). Relative fluorescence recovery at the plateau corresponding to the curves was provided for all markers in Fig. 2h, l, p. Curves in **Extended data 3a, c** were the same set as in Fig. 2c, d. Curves in **Extended data 3f, i** were the same set as in Fig. 2k. Curves in **Extended data 3k, m** were the same set as in Fig. 2o.

**Extended data 4.**
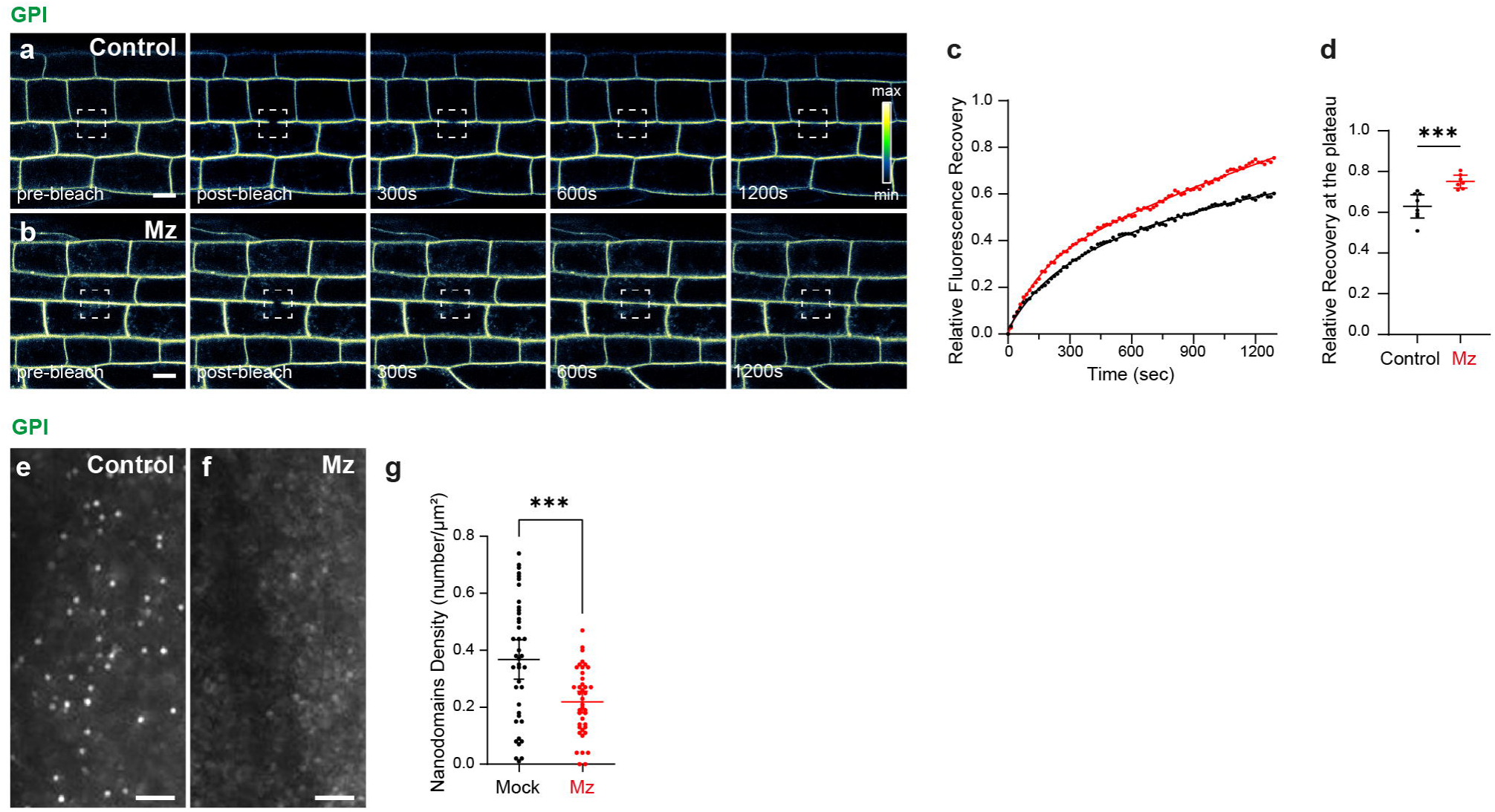
The mobility of a minimal GPI-anchored protein and the formation of GPI nanodomains are VLCFA-dependent. (**a, b**) Fluorescence Recovery After Photobleaching (FRAP) experiments in root epidermal cells of plants expressing the outer leaflet minimal GPI-GFP protein in either control condition (**a**) or upon Mz treatment (**b**). (**c**) Relative fluorescence recovery curves corresponding to **a, b**. (**d**) Relative fluorescence recovery at the plateau corresponding to the curves in **c**. As compared to the control condition, GPI displayed an enhanced mobility when plants were treated with Mz (n=7-8). **(e, f)** Total Internal Reflection Fluorescence (TIRF) microscopy images of GPI-GFP acquired at the PM surface of root epidermal cells. (**e**) GPI constitutive nanodomains are disrupted in Mz treatment (**f**). (**g**) Quantification of the nanodomain density (number / µm²) corresponding to **e, f** (n=40). Statistical tests were performed by ANOVA Kruskal-Wallis in **c** and by Mann-Whitney in **g**, ns *P* >0.05, * *P* <0.05, ** *P* <0.01, *** *P* <0.001. Scale bars are 10 µm in **a, b** and 2 µm in **e, f**.

**Extended data 5.**
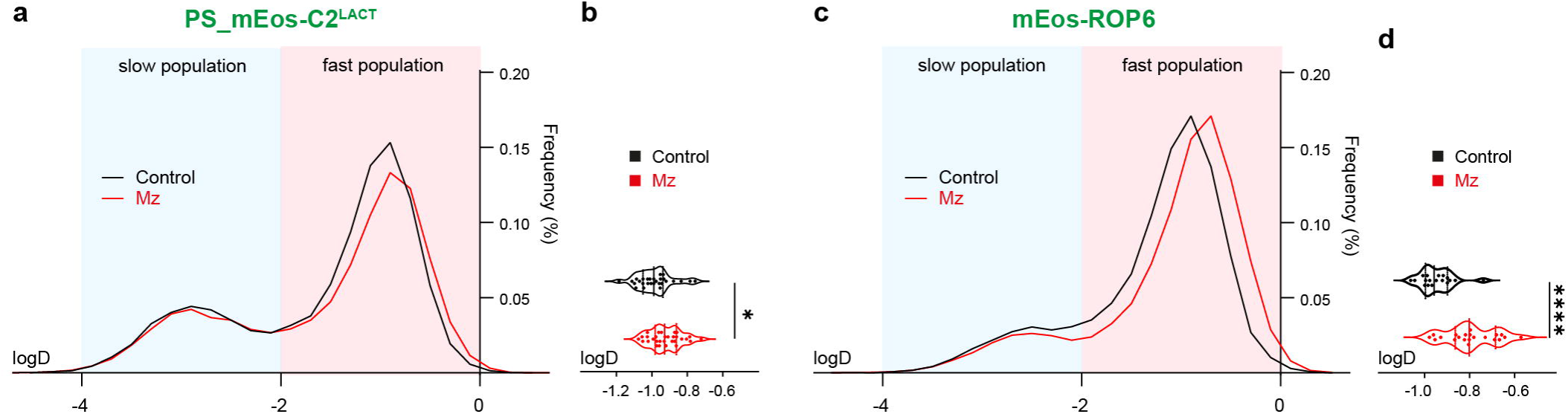
Single Particle Tracking (SPT) PhotoActivated Localization Microscopy (PALM) of the PS biosensor mEos-C2^LACT^ and mEos-ROP6. Both the PS biosensor mEos-C2^LACT^ (**a, b**) and mEos-ROP6 (**c, d**) show a significant increase of mobility upon Mz treatment (n=28-30 in **b**, n=18-21 in **d**). Statistical tests were performed by t-test, ns *P* >0.05, * *P* <0.05, ** *P* <0.01, *** *P* <0.001, **** *P* <0.0001.

**Extended data 6.**
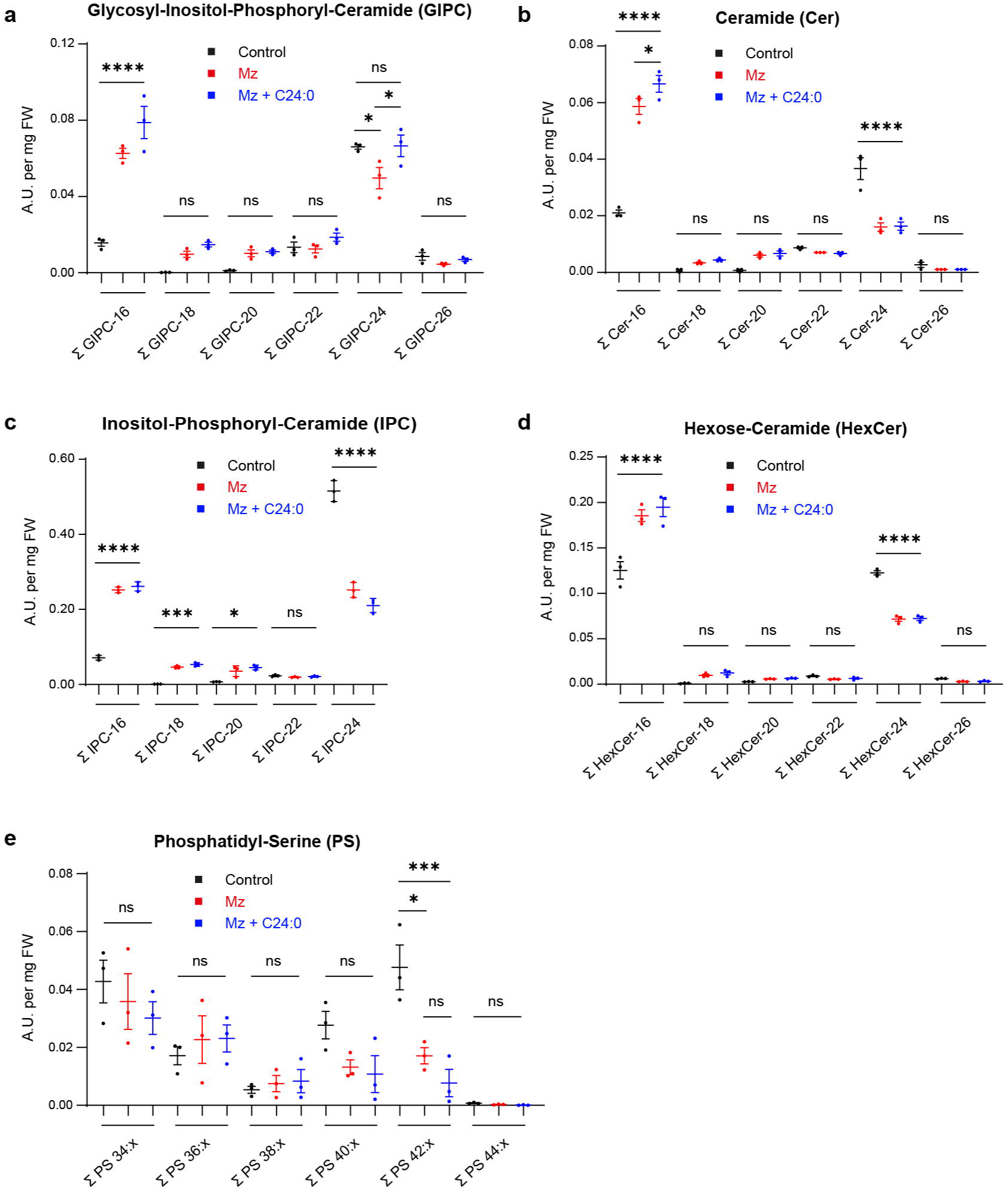
Lipidomic analysis of extracts from whole seedlings supporting Fig. 6. LC-MS/MS analyses of lipid extracts from whole seedlings in control condition (black), treated with Mz for 24h (red) or treated with Mz and 24:0 fatty acids for 24h (blue). (**a**) 24;0 fatty acids rescued VLCFA in the final and main pool of sphingolipid, *i.e* the Glycosyl-Inositol-Phosphoryl-Ceramides (GIPC) (n=3) but not in either the biosynthetic intermediate pools of sphingolipids (such as Ceramides (Cer, **b**) and Inositol-Phosphoryl-Ceramide (IPC, **c**)), Hexose-Ceramides pool (Hex-Cer, **d**) or phosphatidylserine pool (PS, **e**) pool (n=3). Statistical tests were performed by ANOVA, ns *P* >0.05, * *P* <0.05, ** *P* <0.01, *** *P* <0.001, **** *P* <0.0001.

